# Surfactin production is not essential for pellicle and root-associated biofilm development of *Bacillus subtilis*

**DOI:** 10.1101/865345

**Authors:** Maude Thérien, Heiko T. Kiesewalter, Emile Auria, Vincent Charron-Lamoureux, Mario Wibowo, Gergely Maróti, Ákos T. Kovács, Pascale B. Beauregard

**Affiliations:** Centre SÈVE, Département de biologie, Faculté des Sciences, Université de Sherbrooke, Sherbrooke, Canada; Bacterial Interactions and Evolution Group, DTU Bioengineering, Technical University of Denmark, Kgs Lyngby, Denmark; Biology department, Ecole Normale Supérieure Paris-Saclay, Paris-Saclay University, Cachan, France; Natural Product Discovery Group, DTU Bioengineering, Technical University of Denmark, Kgs Lyngby, Denmark; Institute of Plant Biology, Biological Research Center of the Hungarian Academy of Sciences, Szeged, Hungary

**Keywords:** *Bacillus subtilis*, biofilm, surfactin, plant root colonization, pellicle

## Abstract

Secondary metabolites have an important impact on the biocontrol potential of soil-derived microbes. In addition, various microbe-produced chemicals have been suggested to impact the development and phenotypic differentiation of bacteria, including biofilms. The non-ribosomal synthesized lipopeptide of *Bacillus subtilis*, surfactin, has been described to impact the plant promoting capacity of the bacterium. Here, we investigated the impact of surfactin production on biofilm formation of *B. subtilis* using the laboratory model systems; pellicle formation at the air-medium interface and architecturally complex colony development, in addition to plant root-associated biofilms. We found that the production of surfactin by *B. subtilis* is not essential for pellicle biofilm formation neither in the well-studied strain, NCIB 3610, nor in the newly isolated environmental strains, but lack of surfactin reduces colony expansion. Further, plant root colonization was comparable both in the presence or absence of surfactin synthesis. Our results suggest that surfactin-related biocontrol and plant promotion in *B. subtilis* strains are independent of biofilm formation.

## 1. Introduction

Several species from the “*Bacillus subtilis* complex” are well-characterized plant growth-promoting rhizobacteria (PGPRs), providing various beneficial activities for plants and inhibiting fungal and bacterial pathogens [1]. Many strains of *Bacillus subtilis, Bacillus amyloliquefaciens* and *Bacillus velezensis* are currently used in organic and traditional agriculture to prevent infection and/or increase yields of various crops [2–4]. These species are of particular interest because they can form stress-resistant endospores, a cell-type ideal for product formulation. Most PGPR *Bacillus spp.* also produce a wide range of bioactive molecules, such as lipopeptides, which directly influences plant growth and defence [5].

Many of these molecules are synthesized by multienzyme-complexes called non-ribosomal peptide synthetases (NRPS) [6]. *B. subtilis* NCIB3610 possesses 3 NRPS clusters and one NRPS/polyketide synthetase (PKS) cluster, which is few compared to the bioactive molecule synthesis capacity of *B. velezensis* strains [1]. Bacillaene, a broad spectrum antibiotic, is synthesized by proteins encoded in 80 kB *pksA-S* cluster [7]. The *ppsA-E* encodes for the peptide synthetase responsible for the synthesis of plipastatin (fengycin family), a strong antifungal molecule [5,8], while the siderophore bacillibactin is synthesized by the product of the *dhbA-F* operon [9]. Finally, SrfAA-AD produces versatile molecules from the surfactin family [10].

Surfactin molecules are composed of a heptapeptide, i.e. two acidic and five nonpolar amino acids, interlinked with a β-hydroxy fatty acid, and condensed in a cyclic lactone right structure [10,11]. The amino acid sequence, the length, and the branching of the fatty acid moiety can vary in surfactin molecules produced by different *Bacillus* species, strains and/or growth conditions [12]. For example, on tomato roots *B. amyloliquefaciens* S499 produces surfactin variants with C_12_, C_13_, C_14_ and C_15_ acyl chains, the last two composing more than 80% of total surfactins produced in these conditions [13]. Surfactin, as its name suggests, is an extremely powerful biosurfactant, and thus helps bacteria moving on solid surface [6,14–18]. These molecules are abundantly produced when *B. subtilis* colonizes plant roots, and they elicit the induce systemic resistance in plants [19–22].

A strong link between biofilm formation and surfactin production was suggested for different *Bacillus* species. Mutations in the surfactin synthesis operon were reported to cause partial to severe biofilm defect *B. velezensis* FZB42 and *B. amyloliquefaciens* UMAF6614 [23,24]. Non-surfactin producer strains of UMAF6614 were also impaired in root colonization [25]. Under specific laboratory growth conditions (i.e. exponentially growing cells inoculated into lysogeny broth medium), surfactin was shown to trigger biofilm formation in *B. subtilis* via a pore-forming activity, which causes intracellular potassium leakage sensed by KinC that in turns activate the genetic pathway responsible for biofilm formation [26]. This “quorum-sensing like” activity was demonstrated in the model strain NCIB3610. Similarly to reports in *B. velezensis* and *B. amyloliquefaciens*, a surfactin deletion mutant of *B. subtilis* 6051 was shown to be defective for biofilm formation and root colonization [19]. However, a different study showed that biofilms formed by *B. subtilis* tomato rhizoplane isolates had comparable dry weight among wild-type and surfactin mutants [27]. Finally, deletion of *sfp*, which is known to be involved in the production of surfactin since it encodes for a 4′ phosphopantetheinyl transferase that activates the peptidyl carrier protein domains from the NRPS machinery, impairs biofilm formation in *B. subtilis* 3610 [28]. Since *sfp* mutation is defective for the synthesis of all NRP-derived molecules (surfactin, bacillibactin, plipastatin and bacillaene), this impair in biofilm formation could be due to a defect in other biosynthetic pathways than surfactin [14]. These conflicting reports and recent results in our laboratories lead us to revisit the importance of surfactin for biofilm formation of *B. subtilis in vitro* and *on planta*.

## 2. Material and Methods

### 2.1 Strains, media, and chemicals

Strains used in the study are listed in Table S1. For routine growth, cells were propagated on lysogeny broth (LB; Luria-Bertani or Lenox broth) medium. When necessary, antibiotics were used at the following concentrations: MLS (1 μg mL^−1^ erythromycin, 25 μg mL^−1^ lincomycin); spectinomycin (100 μg mL^−1^); chloramphenicol (5 μg mL^−1^) and kanamycin (10 μg mL^−1^). New *B. subtilis* isolates were obtained from 5 sampling sites in Germany and Denmark (see Table S1 for coordinates) by selecting for spore formers in the soil. Soil samples were mixed with 0.9% saline solution, vortexed on a rotary shaker for 2 min, incubated at 80°C for 25 min and serially diluted on 2×SG medium solidified with 1.5% agar [29]. Highly structured colonies were targeted and isolation of *B. subtilis* strains was confirmed using 16S sequencing followed by whole genome [30]. New isolates and their *srfAC::spec* derivatives (*srfAC::spec* marker was transferred from DS1122 [31]) were labeled with constitutively expressed *gfp* from P_hyperspank_ using phyGFP plasmids that integrates into the *amyE* locus [32].

**Table S1.**
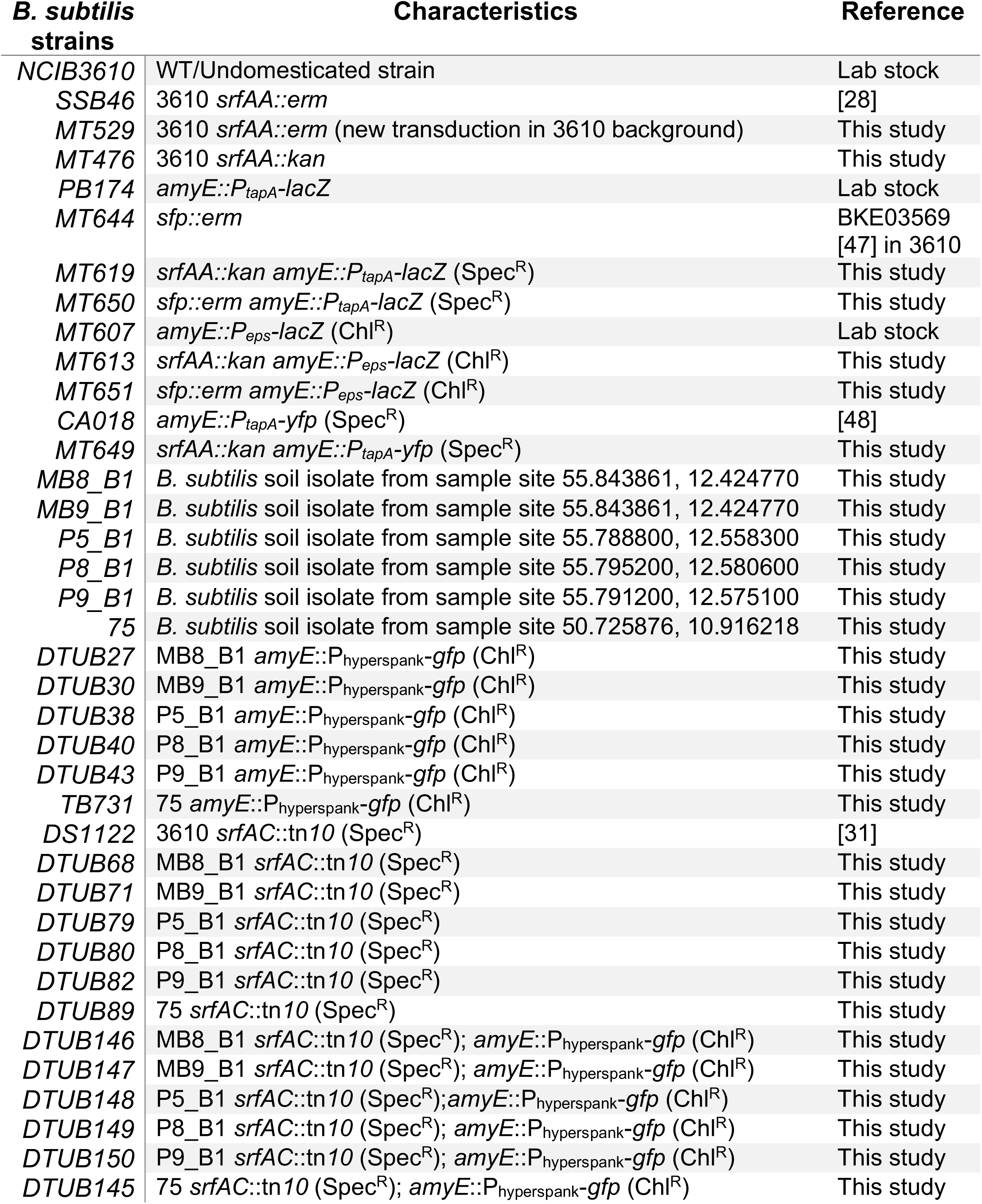
Strains used in this study.

All solvents used for HRMS and chromatography were VWR Chemicals LC-MS grade, while for metabolites extraction the solvents were HPLC grade (VWR Chemicals). Surfactin standard was purchased from Sigma-Aldrich (Cat. No. S3523).

### 2.2 Culture conditions

For pellicles assays, cells were pre-grown for 2 hours and diluted 1:100 in 3mL LB 3 times, and at the last incubation cells were grown until they reach an OD_600_ between 0.3 and 0.6. OD_600_ was then adjusted at 0.3 with LB, and 13.5 μL were used to inoculate 1mL of medium in a 24-well plates. Media used for these experiments were MSgg [28] and MSNc + Pectin (MSN: 5mM Potassium phosphate buffer pH7, 0.1M Mops pH7, 2mM MgCl_2_, 0.05mM MnCl_2_, 1μM ZnCl_2_, 2 μM thiamine, 700 μM CaCl_2_, 0.2% NH_4_Cl; 0,5% cellobiose and 0,5% pectin (Sigma)). Incubation was done at 30°C. For pellicle assays of recent *B. subtilis* soil isolates and its mutant derivatives, three to four colonies were inoculated in 3 ml LB and incubated at 37°C with shaking at 225 rpm for 4 h. The OD_600_ was adjusted to 1.5 and 1 % inoculum of the pre-grown culture was used to seed bacterial biofilms in MSgg [28] or MOLP [33] media at 30°C. For colony biofilms, one colony was inoculated in 3 mL LB and rolled for 3h at 37°C. The culture was adjusted to an OD_600_ of 1, then 2μL were spotted on solidified (1.5% agar) MSgg media.

Col-0 *A. thaliana* ecotype was used throughout the study. In the Canadian laboratory, seeds were surface-sterilized with 70% ethanol followed by 0.3% sodium hypochlorite (v/v) and germinated on Murashige-Skoog medium (Sigma) 0.7% agar with 0.05% glucose in a growth chamber at 25°C. Root colonization assay were performed using MSNg (MSN supplemented with 0.05% glycerol) as described in [34]. In Denmark, *Arabidopsis* seeds were surface sterilized using 2% (v/v) sodium hypochlorite with mixing on an orbital shaker for 20 min and then washed five times with sterile distilled water. The seeds were placed on pre-dried Murashige and Skoog (MS) basal salts mixture (2.2 g l^−1^, Sigma) containing 1% agar in an arrangement of approximately 20 seeds per plate at a minimum distance of 1 cm. After 3 days of incubation at 4°C, plates were placed at an angle of 65° in a plant chamber with a light regime of 16 h light (24°C)/8-h dark (21°C). After 6 days, homogenous seedlings ranging 0.8-1.2 cm in length were selected for root colonization assay. Seedlings were transferred into 48-well plates containing 270 μl of MSNg medium [34] per well. The wells were supplemented with 30 μl of exponentially growing bacterial culture diluted to OD_600_ = 0.2. The sealed plates were incubated at a rotary shaker (90 rpm) at 30°C for 18 h. After the incubation, plants were washed three times with MSNg to remove non-attaching cells and then transferred to a glass slide for imaging using CLSM.

### 2.3 Beta-galactosidase assays

From pellicle biofilm assays, spent medium was cautiously removed from the wells. The pellicle was then collected in 1mL of Z-buffer (40 mM NaHPO_4_; 60 mM Na_2_HPO_4_; 1 mM MgSO_4_; 10 mM KCl) and transferred in a 1.5mL tube. The suspensions were sonicated with 1 second pulses (30% power) for 10 seconds total to break the biofilms, and OD_600_ was measured. Then, 2-mercaptoethanol (final concentration of 38 mM) and freshly prepared lysozyme in Z-buffer (final concentration of 20 µg mL^−1^) were added. Suspensions were incubated for 30 min at 30°C, diluted and 100µL of an ONPG solution (4 mg mL^−1^ in Z-buffer with 38mM of 2-mercaptoethanol) were added. 250µL of Na_2_CO_3_ 1M were added when solutions started turning yellow, and the reaction time was recorded. The A_420nm_ and OD_550nm_ were measured for each solution, and the Miller Units were calculated using: Miller Units = 1000 × [(A_420nm_ − 1.75 × OD_550nm_)] / (T_min_ × V_ml_ × OD_600_)

### 2.4 Microscopy

To visualize bacteria on root surfaces in the Canadian laboratory, seedlings were examined with a Zeiss Axio Observer Z1 microscope equipped with a 20X/0.8 Plan-Apochromat objective, and whole root pictures were taken with a Zeiss Axiocam 506 mono. Figure 3A presents representative images of the various mutant and time of colonization. The fluorescence signal was detected using a YFP filter (ex: 500/20, em: 535/30) and a CFP filter for autofluorescence of the root (ex: 436/20, em: 480/40). All images were taken at the same exposure time, processed identically for compared image sets, and prepared for presentation using Zeiss Zen 2.0 software. Each image is representative of at least 12 root colonization assays performed in three independent experiments. Quantification was performed using CellProfiler 3.0 (cellprofiler.org) [35].

**Figure 1.**
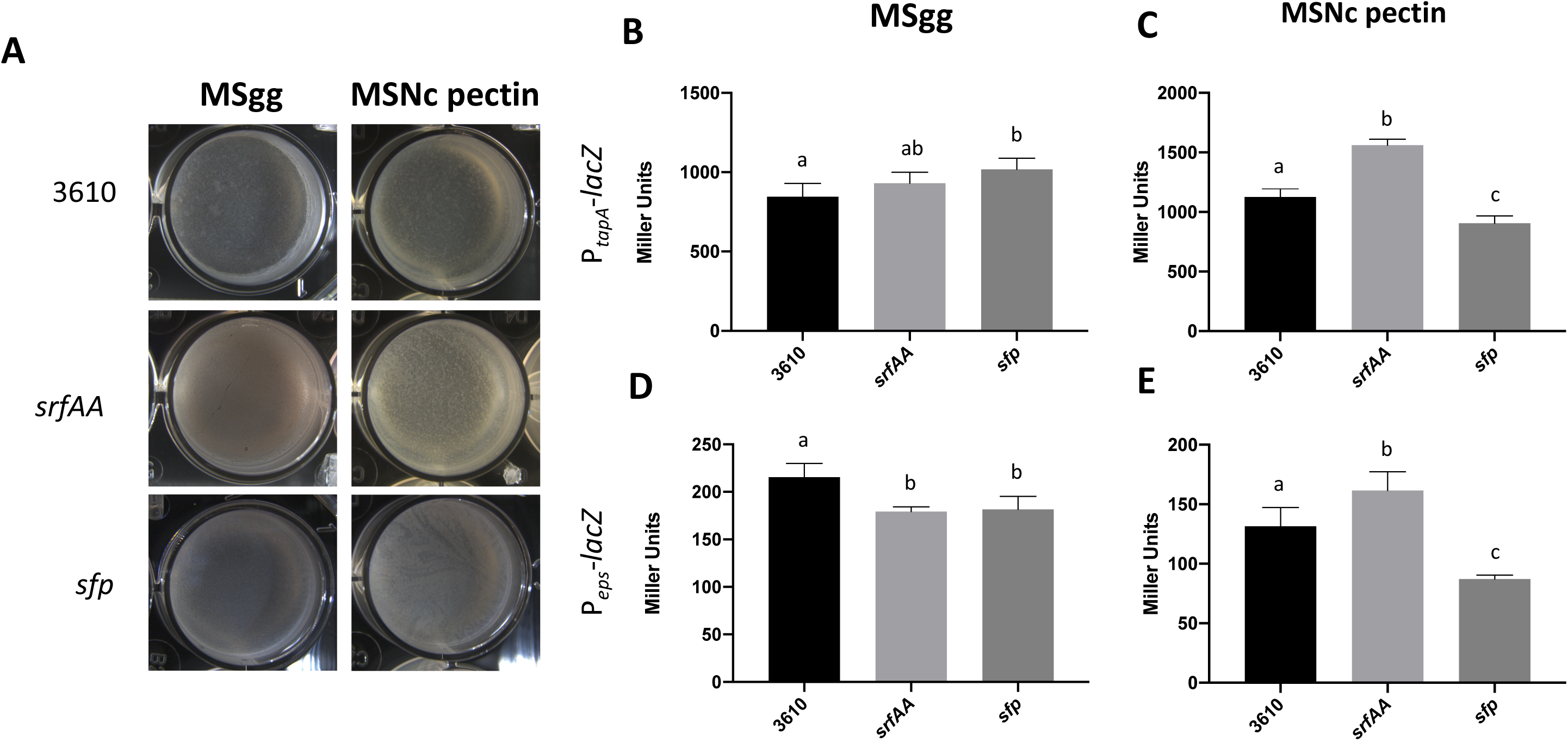
Pellicle formation is mildly affected by *srfAA* or *sfp* deletion. (A) Top-down view of pellicle assay in which the indicated mutants were incubated for 24 h at 30 °C in MSgg or in MSNc + pectin. Results are representative of three experiments. (B-D) β-galactosidase activities of WT (3610), *srfAA* or *sfp* mutant harbouring the P_*tapA*_-lacZreporter (B and C) or the P_*eps*_-lacZ reporter (D and E). Cells were grown in standing MSgg (B and D) or MSNc + pectin (C and E) pellicles for 20 hours. Values represent the mean of five technical replicates, and the experiments are representative of at least three independent biological replicates. Error bars represent standard deviation, and letters represent = P <0.05.

**Figure 2.**
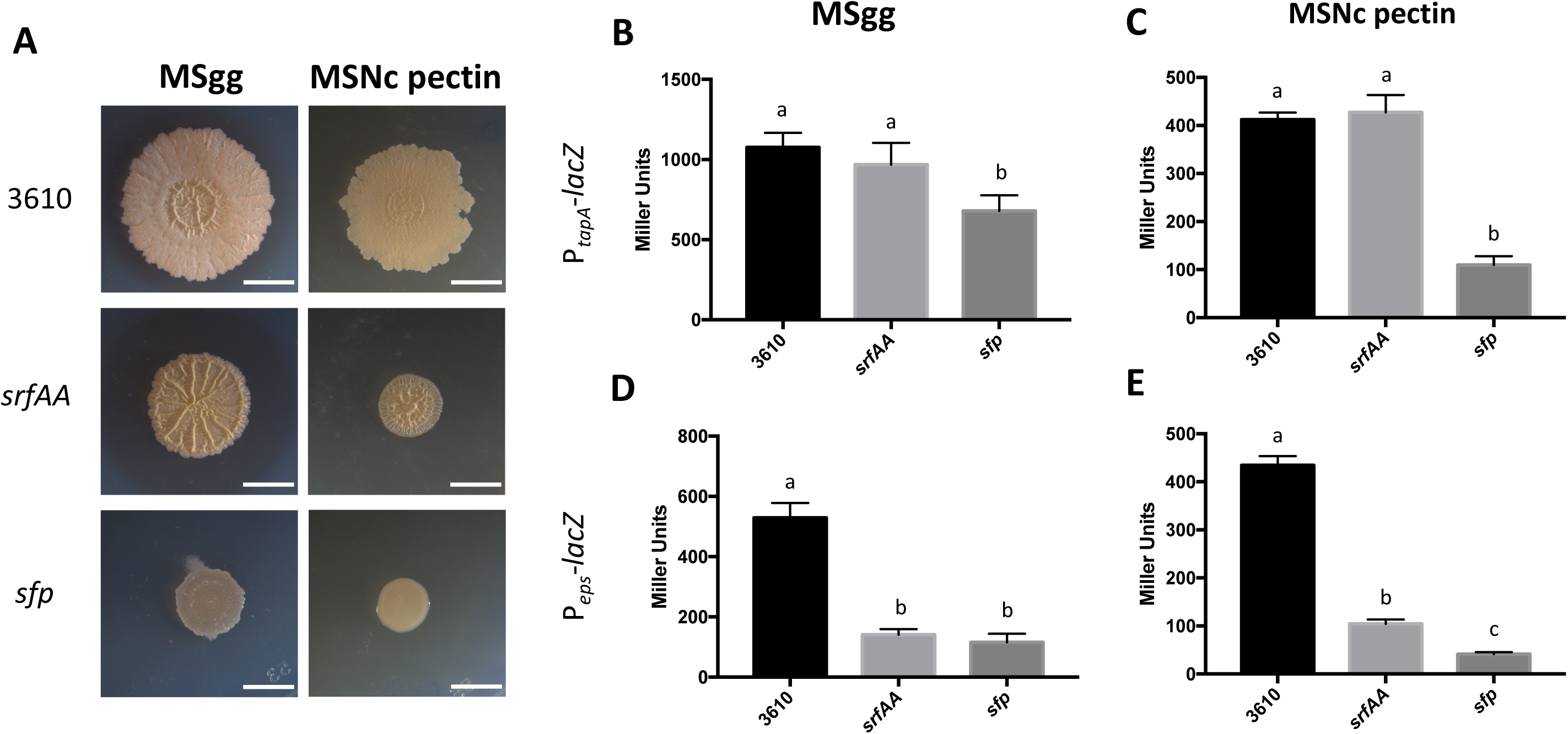
*srfAA* or *sfp* influence colony formation. (A) Top-down view of colonies incubated for 72 h at 30 °C on solid MSgg or MSNc + pectin. Results are representative of three experiments. Scale bar are 5 mm. (B-D) β-galactosidase activities of WT (3610), *srfAA* or *sfp* mutant harbouring the P_*tapA*_-*lacZ* reporter (B and C) or the P_*eps*_-*lacZ* reporter (D and E). Cells were grown on solid MSgg (B and D) or MSNc + pectin (C and E) for 20 hours. Values represent the mean of six technical replicates, and the experiments are representative of at least three independent biological replicates. Error bars represent standard deviation, and letters represent = P <0.05.

**Figure 3.**
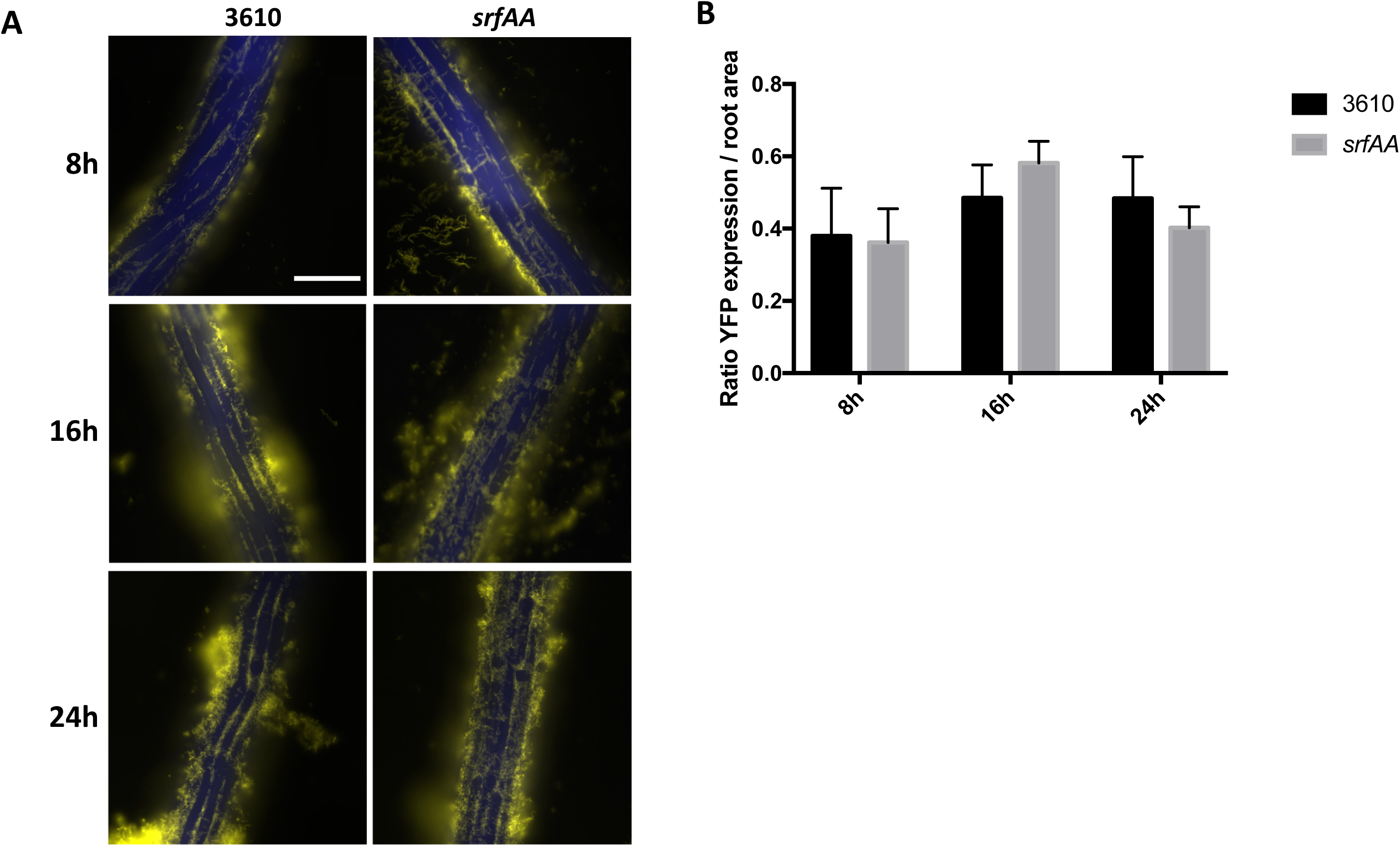
Surfactin is not required for root colonization. (A) 3610 cells harbouring P_*tapA*_-*yfp* co-incubated with *A. thaliana* seedlings and imaged at 8, 16 and 24 h post-inoculation. Shown are overlays of fluorescence (false-colored green for YFP, and blue for CFP filter - which represents the autofluorescence of roots). Pictures are representative of 12 independent roots. Scale bar is 100 μm for all images. (B) The entire root was imaged at 20×, and numbers of fluorescent pixels was counted and then divided by the root’s area (also measured in pixel), allowing quantification of biofilm-forming cells present on the root. For each strain, the bar represents the mean and standard deviation of at least four technical replicates; experiment is representative of three independent biological replicates. There was no statistical difference between 3610 and *srfAA* in the various conditions.

In Denmark, the washed plant roots were transferred to microscope slides and gently sealed with cover slips. Plant root colonization was analysed with a confocal laser scanning microscope (TCS SP8 (Leica) equipped with an argon laser and a Plan-Apochromat 63x/1.4 Oil objective). Fluorescent reporter excitation was performed at 488 nm for green fluorescence, while the emitted fluorescence was recorded at 520/23 nm. Single-layer images were acquired and processed with the software ImageJ (National Institutes of Health). Each image is representative of 2 root colonization assays performed in two independent experiments.

### 2.5 Chemical extraction of secondary metabolites from bacterial cultures

Bacterial strains were cultured on MSgg agar plates for 3 days at 30 °C. An agar plug (6 mm diameter) of each bacterial cultures was transferred to a vial and extracted with 1 mL of isopropanol:ethyl acetate (1:3, v/v) with 1% formic acid. The vials were placed in an ultrasonic bath at full effect for 60 min. Extracts were then transferred to new vials, evaporated to dryness under N_2_, and re-dissolved in 300 µL of methanol for further sonication over 15 min. After centrifugation at 13400 rpm for 3 min, the supernatants were transferred to new vials and subjected to ultrahigh-performance liquid chromatography-high resolution mass spectrometry (UHPLC-HRMS) analysis.

### 2.6 UHPLC-HRMS analysis

UHPLC-HRMS was performed on an Agilent Infinity 1290 UHPLC system equipped with a diode array detector. UV–visible spectra were recorded from 190 to 640 nm. Liquid chromatography of 1 µL extract was performed using an Agilent Poroshell 120 phenyl-hexyl column (2.1 × 150 mm, 2.7 μm) at 60 °C with acetonitrile and H_2_O, both buffered with 20 mM formic acid, as mobile phases. Initially, a linear gradient of 10% acetonitrile in H_2_O to 100% acetonitrile over 15 min was employed, followed by isocratic elution of 100% acetonitrile for 2 min. The gradient was returned to 10% acetonitrile in H_2_O in 0.1 min, and finally isocratic condition of 10% acetonitrile in H_2_O for 2.9 min, all at a flow rate of 0.35 mL/min. MS detection was performed in positive ionization on an Agilent 6545 QTOF MS equipped with an Agilent Dual Jet Stream electrospray ion source with a drying gas temperature of 250 °C, drying gas flow of 8 L/min, sheath gas temperature of 300 °C, and sheath gasflow of 12 L/min. Capillary voltage was set to 4000 V and nozzle voltage to 500 V. MS data processing and analysis were performed using Agilent MassHunter Qualitative Analysis B.07.00.

### 2.7 Genome re-sequencing

Genomic DNA of 3610, SSB46 and MT529 were isolated using Bacterial and Yeast Genomic DNA kit (EURx). Re-sequencing was performed on an Illumina NextSeq instrument using V2 sequencing chemistry (2×150 nt). Base-calling was carried out with “bcl2fastq” software (v.2.17.1.14, Illumina). Paired-end reads were further analyzed in CLC Genomics Workbench Tool 9.5.1. Reads were quality-trimmed using an error probability of 0.05 (Q13) as the threshold. Reads that displayed ≥80% similarity to the reference over ≥80% of their read lengths were used in mapping. Quality-based SNP and small In/Del variant calling was carried out requiring ≥10× read coverage with ≥25% variant frequency. Only variants supported by good quality bases (Q ≥ 30) on both strands were considered.

## 3. Results

### 3.1 Absence of surfactin has no effect on pellicle formation of NCIB 3610

To assess the importance of surfactin production for biofilm development, pellicle formation, a biofilm on the air-medium interface was first examine in liquid biofilm-inducing medium, i.e. MSgg and MSNc + pectin. MSgg induces biofilm formation via iron availability and glutamate, while pectin, a plant-derived polysaccharide, is the main environmental cue inducing biofilm formation in MSNc + pectin [34]. Since both media present different cues for the bacterial cells, and that pectin was shown to strongly induce surfactin production [36], importance of this molecule for biofilm formation could vary according to the medium used. As shown in Fig. 1A, deletion of *srfAA*, and consequently absence of surfactin, does not visibly affect pellicle formation in either liquid media. Similarly, a strain deleted for *sfp*, which is defective for synthesis of all NRP-derived molecules, is also able to form pellicle in both media.

Importantly, the 3610 *srfAA* deletion strain used here was newly created (harbouring a kanamycin resistant gene) and did not match the pellicle formation phenotype of the originally published laboratory stock, 3610 *srfAA::erm* (SSB46; [28]), the latter showing an important delay in pellicle formation (Fig. S1). When the *srfAA::erm* marker was re-introduced into 3610 by SPP1 phage transduction, the newly obtained *srfAA::erm* strain (MT529) displayed comparable pellicle development to 3610 and *srfAA::kan* strains. Consequently, the genomes of 3610, SSB46, and the newly created MT529 strains were re-sequenced. In addition to the *srfAA::erm* mutation, SSB46 strain contained six point mutations that did not exist the ancestral 3610 or the re-created MT529 strain (see Table S2). However, deletion mutants of the SNP harbouring genes combined with *srfAA::kan* did not recapitulate the important defect observed with SSB46 strain (Fig. S1), suggesting that the mutation causing the defect is not a loss-of-function or that certain combination of SNPs are responsible for the observed phenotype of SSB46 strain.

**Fig S1.**
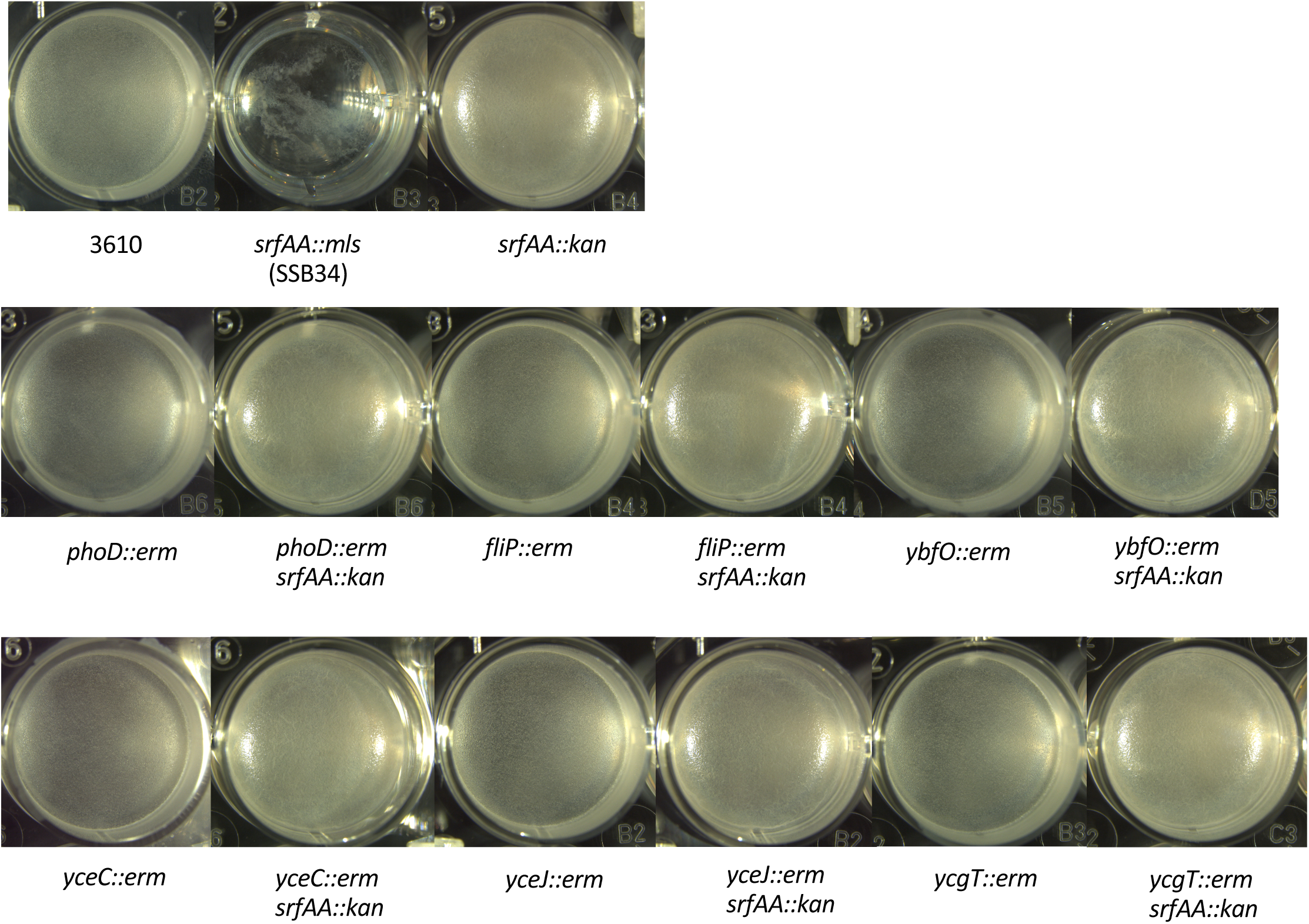
Pellicle formation with deletion mutants identified as containing mutations in strain SSB46. Pellicles were formed in MSgg for 24 h at 30 °C. Pictures are representative of biological duplicates.

Importance of surfactin and *sfp* for activation of the two main operons involved in biosynthesis of the extracellular matrix, i.e. *tapA-sipW-tasA* and *epsA-O*, was further examined using transcriptional *lacZ* fusions. As shown in Fig. 1B and D, absence of surfactin or deletion of *sfp* have little to no effect on *tapA* transcription, and slightly decreases *epsA-O* transcription in MSgg. In MSNc pectin, absence of surfactin actually increases *tapA* and *epsA-O* transcription (Fig. 1C), which also correlates with the more vigorous aspect of pellicles (see Fig.1A). In the same medium, absence of *sfp* impairs transcription of both biofilm operons. In summary, in liquid media *srfAA* or *sfp* deletion has only mild impacts on pellicle biofilm formation in *B. subtilis* 3610.

### 3.2 Deletion of sfp and srfAA alters colony structure

Biofilm strength can also be evaluated using the complex architecture of colony biofilms growing on solid biofilm-inducing media. Since surfactin is a biosurfactant [17,37], its absence might have more severe effect on solid media as observed for sliding, a matrix dependent colony expansion [17]. Indeed, as shown in Fig. 2A, colonies of *srfAA* show less spreading on solid MSgg and MSNc pectin, but are still very wrinkly. These wrinkles are likely composed mostly of proteinaceous (*TasA*) fibres, since expression of P_*eps*_ is drastically reduced by absence of surfactin, while P_*tapA*_ is not affected (Fig. 2B). Interestingly, the *sfp* mutant produces small, flat colonies in both media. This strain also has significantly reduced LacZ activity for both biofilm reporters (P_*tapA*_ and P_*eps*_) and media, which correspond to the flat phenotype of the colonies. While calculation of the miller units includes normalization for cell number (OD_600_), this lack of biofilm gene expression could also be attributable to lack of cell growth and incapacity to reach the cell density required for biofilm formation.

**Fig S2.**
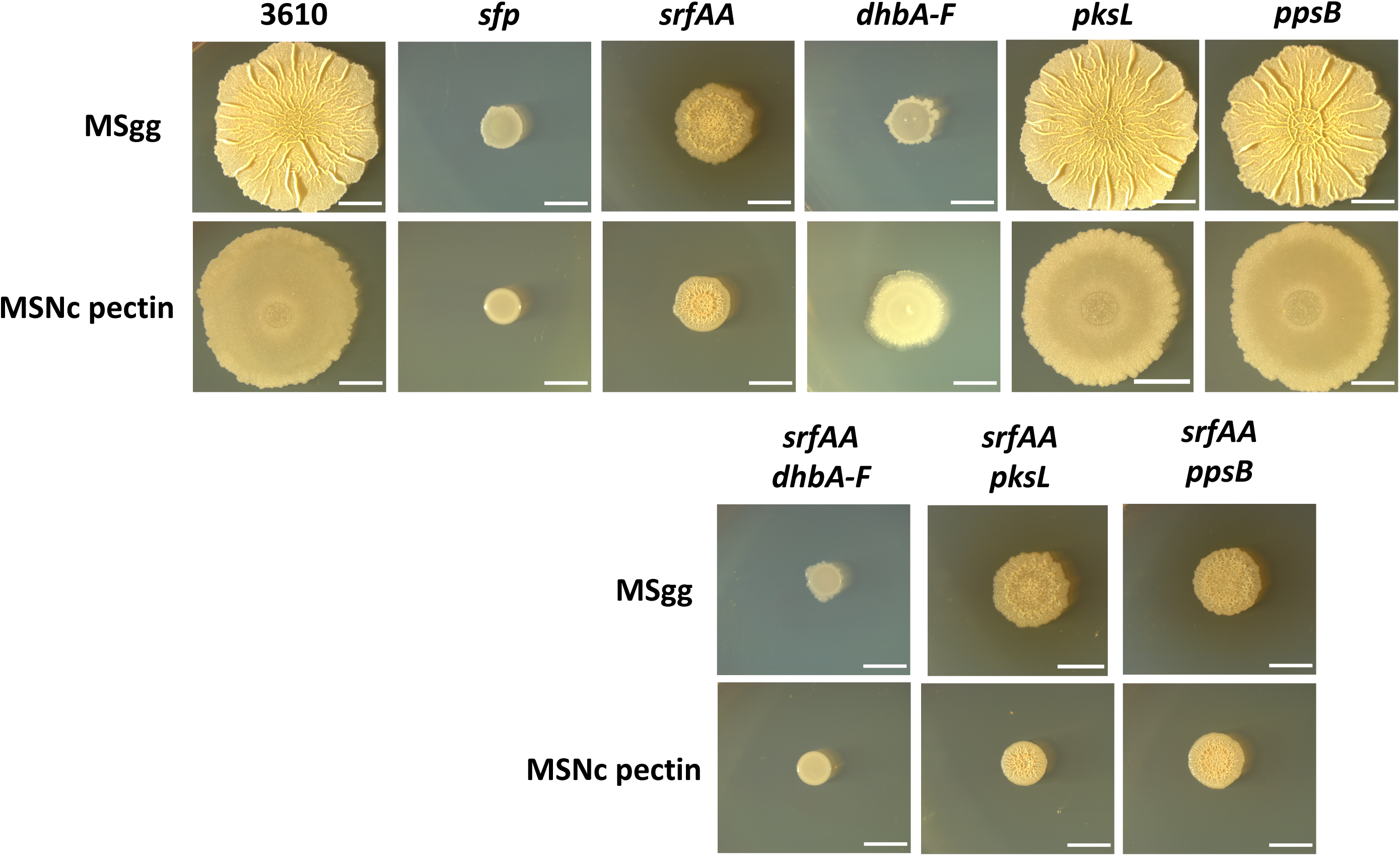
Impact of NRPs mutations on colony complexity. Top-down view of colonies incubated for 72 h at 30 °C on solid MSgg or MSNc + pectin. Results are representative of at least two experiments. Scale bars are 5 mm.

Colony morphology results on solid media clearly show that *srfAA* and *sfp* deletion lead to phenotypes, indicating that in the latter absence of other molecule(s) synthesized via NRP machinery also impacts biofilm formation. Thus, we examined deletion mutants for bacillibactin (*dhbA-F*), plipastatin (*ppsB*) and bacillaene (*pksL*) (Fig. S2). The mutant defective for *B. subtilis* siderophore bacillibactin showed small, almost featureless colonies on both media, suggesting an important role for iron-acquisition molecules in this process. The double *srfAA dhbA-F* deletion recapitulated the *sfp* phenotype, suggesting that on solid media, both molecules are required for robust biofilm formation.

### 3.3 *Surfactin is not required for root colonization by* B. subtilis

In a natural environment, surfactin production is triggered by contact with plant roots few hours before biofilm formation [36]. Thus, we evaluated the importance of surfactin for root colonization of *A. thaliana* seedlings, using the system described in [34]. *B. subtilis* root colonization was monitored using a P_*tapA*_-*yfp* reporter, allowing us to identify cells actively forming a biofilm on roots. Since absence of surfactin might only delay, instead of inhibit, root colonization, different time points after inoculation were examined. As shown in Fig. 3A, there was no apparent difference in the root colonization patterns and capacities of WT and *srfAA* cells. We validated these observations by imaging whole roots and determining the ratio of YFP expression/root area, which gives us a quantitative measurement of colonization. Indeed, while colonization somewhat varied from one seedling to another, overall there was no significant difference between WT and *srfAA* root colonization at any time points.

**Table S2.** Point mutations detected only in SSB46 (*srfAA::erm*) Positions are indicated in reference to the genome of 3610 (NCBI Gene Bank sequence CP020102.1)

| Position | Coding region change | Amino-acid change | Gene |
| --- | --- | --- | --- |
| 251310 | deletion of CT and insertion of TA | Leu430Tyr | *ybfO* |
| 284231 | A to C | Lys59Asn | *phoD* |
| 312530 | Insertion of G | Glu110fs | *yceC* |
| 319949 | C to T |  | *yceJ* |
| 353100 | deletion of AGCAGCTGATCG | Ile70_Leu73del | *ycgT* |
| 1705268 | G to T |  | *fliP* |

### 3.4 *Surfactin is dispensable for pellicle and plant-associated biofilm formation in recent soil* B. subtilis *isolates*

To address the generality of lack of surfactin production on pellicle formation ability, we tested pellicle biofilm development of 6 newly isolated *B. subtilis* strains recovered from soil samples. As the essentiality of surfactin production for pellicle development has been demonstrated on MOLP medium for *B. amyloliquefaciens* (previously identified as *B. subtilis*) UMAF6614 [24], pellicle formation was followed both on MSgg and MOLP liquid media that revealed no observable difference between wild-type and their surfactin mutant derivatives (Fig. 4). Additionally, plant colonization was indistinguishable between the wild-type and *srfAC::spec* strains (Fig. 4). Finally, to demonstrate the surfactin production ability of these new *B. subtilis* strains, the isolates were inoculated to MSgg medium and UHPLC-HRMS analysis was performed on isopropanol:ethyl acetate extracts of the agar medium below the colonies. Chemical analysis of the extract along with a standard demonstrated that each and every isolates produced surfactin, but not their *srfAC::spec* derivatives (Fig. S3).

**Fig 4.**
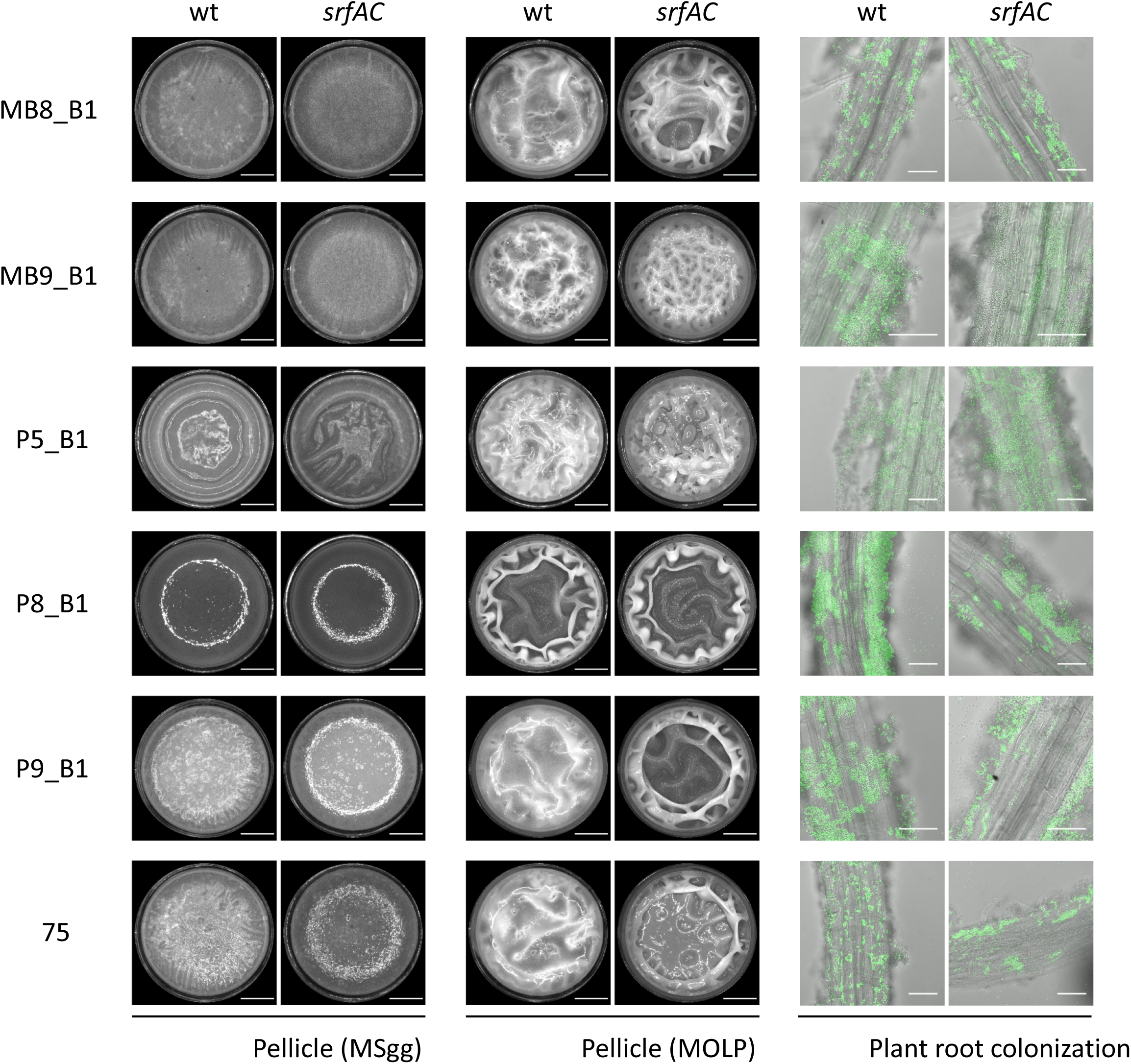
Pellicle (in MSgg and MOLP media) and root associated biofilms of newly isolated *B. subtilis* strains (wt columns) and their respective surfactin mutants (*srfAC* columns). Scale bars indicate 4 mm and 50 μm for pellicle and root colonization images, respectively. Pellicle assays were performed with non-labeled strains, while plant colonization was followed using constitutively expressed GFP from P_*hyperspank*_.

## 4. Discussion

The promiscuous role of secondary metabolites to function as info chemicals has been previously proposed [38–40]. The *B. subtilis* produced surfactin has been reported to lead to induction of biofilm development under non-biofilm inducing conditions [26]. Our results highlight that under biofilm inducing conditions, on liquid or solid biofilm-promoting MSgg and MSNg media, biofilm development of *B. subtilis* 3610 and other newly isolated strains does not require surfactin. Production of both matrix components actually appeared more efficient in MSNc pectin in absence of surfactin, which could be due to the metabolic burden of producing an important NRPs [41,42]. In solid media, *srfAA* and *sfp* mutants display strikingly different phenotypes than WT. In both cases, colony diameter is smaller, stressing the need for surfactin to disperse on a surface [17,18]. Intriguingly, absence of surfactin had a stronger impact on *eps* than on *tapA* transcription, suggesting that surfactin and/or colony spreading might be involved in regulating exopolysaccharides production on solid surface. This regulation would be independent from SinR and AbrB, which act identically on both operons [43].

**Fig S3.**
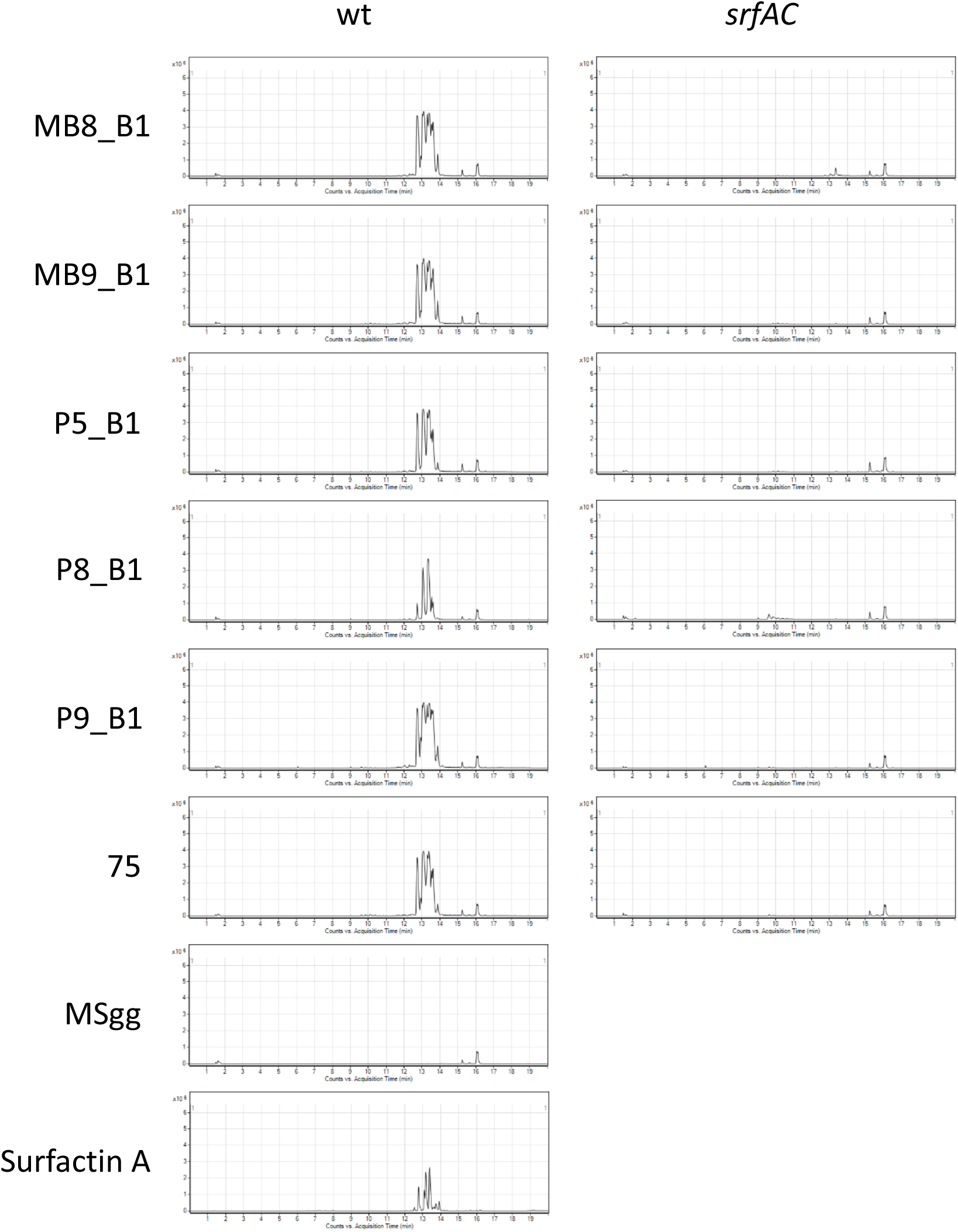
Extracted ion chromatograms (m/z 1000-2000) showing the presence of surfactin produced by the newly isolated *B. subtilis* strains grown on MSgg agar medium and the lack of surfactin production in the *srfAC* derivates. The chromatograms of the MSgg medium and the surfactin standard are shown below. Surfactins, iturins and fengycins are all in the m/z range 1000–2000 that can be detected by ESI–MS [45,46].

Similarly to pellicle biofilm formation in liquid media, various *srfA* mutants colonize plant roots with an efficacy identical to WT cells. Surfactin production is stimulated by plant polysaccharides such as pectin, as is biofilm formation [34,36]. Thus, our observations suggest that while surfactin production precedes biofilm formation upon contact between cells and roots, both processes are somewhat independent. They also would have independent roles, biofilms favouring root attachment and surfactin production, triggering the induced systemic resistance.

Our results show that for *B. subtilis*, surfactin production is not required for robust biofilm formation, which is in contradiction with many reports for surfactin requirement in various *Bacilli* [23,24,44]. In many of these reports however, the species or the strain examined also produce an iturin, bacillomycin, which is not the case for *B. subtilis* 3610 or the newly isolated *B. subtilis* strains [30]. Of note, Luo *et al.* showed that a *srf* mutant of *Bacillus spp.* 916 produces weaker pellicles in liquid medium, and flat colonies on solid MSgg. However, in this case deletion of *srf* also strongly impairs production of bacillomycin L, which is also required for strong biofilm establishment and rice leaves colonization by *Bacillus spp. 916* [44]. Thus, requirement of surfactin for biofilm formation and plant colonization is likely species- or strain-specific in *Bacillus*, and might depend on the presence of iturin production in these strains. Nevertheless, the importance of surfactin production by PGPR strains of *Bacilli* is primarily for the anti-microbial potential and systemic resistance induction by this multi-functional secondary metabolite.

## Acknowledgement

This project was supported by the Danish National Research Foundation (DNRF137) for the Center for Microbial Secondary Metabolites (to H.T.K, M.W., and Á.T.K.), and by a Discovery Grant and Early Career Researcher Supplement from the Natural Sciences and Engineering Council of Canada (NSERC RGPIN-2014-04628 to P.B.B.).

